# Improving Stereochemical Limitations in Protein-Ligand Complex Structure Prediction

**DOI:** 10.1101/2025.03.25.645362

**Authors:** Ryuichiro Ishitani, Yoshitaka Moriwaki

## Abstract

AlphaFold3 has revolutionized biology by enabling the prediction of protein complexes with various biomolecules, including small molecular ligands. However, the ligand structures predicted by the model often contain stereochemical errors. In this study, our comprehensive evaluation of AlphaFold3 and its clone model, Boltz-1, demonstrated significant limitations in ligand stereochemistry of their predicted structures, including chirality, bond, and angle geometries. To address the issue, we developed the restraint-guided inference method that applies stereochemical restraints during the reverse diffusion process. This approach perfectly reproduces the chirality specified in the input chemical structure and improves bond and angle geometries of ligands, while maintaining comparable performance in binding pose prediction. Our method provides a practical solution to the stereochemical errors in predicting protein-ligand complexes, thereby enhancing applications of structure prediction in structural biology and drug discovery.

## Introduction

The field of protein structure prediction has experienced revolutionary advancements in recent years. Notably, AlphaFold2, introduced by DeepMind in 2021, demonstrated unprecedented accuracy in predicting protein structures^1^. Following this, AlphaFold3 has expanded its capabilities beyond traditional protein structure prediction to include the structural determination of heterogeneous complexes involving RNA, DNA, glycans, and small molecular ligands^2^. These remarkable contributions have received significant recognition within the scientific community and have been awarded the 2024 Nobel Prize in Chemistry. Several derivative models, such as Boltz-1^3^, Protenix^4^, Chai-1^5^, and HelixFold3^6^, have also been developed.

AlphaFold3 employs a diffusion model approach, allowing it to predict structures for various biological molecules. In particular, concerning the prediction of protein-ligand complexes, the AlphaFold3 paper claims that it can achieve greater accuracy than conventional molecular docking methods^2^. However, several issues remain to be addressed before it can be used as a substitute for conventional molecular docking methods. For example, it has been reported that the predicted ligand structure sometimes contains stereochemistry violations^2^, including errors with chirality, despite the correct reference structure given to the input features. These challenges could pose significant limitations for applications in biological research, including drug design and biosynthesis.

In this study, we first evaluate the structure prediction of protein-ligand complexes by AlphaFold3^2^ and its clone Boltz-1^3^, mainly focusing on the stereochemistry of ligands. Our analysis reveals that both models often failed to reproduce the local conformations and chirality of ligands specified by the input. Then, we propose a novel method to address the issues of chirality and local conformation prediction without retraining the model. By applying this approach, we achieve a 100% success rate in chirality reproduction. This research elucidates the limitations of structure prediction models and provides a practical solution to overcome these problems.

## Methods

### Benchmark dataset

We evaluated the accuracy of protein-ligand complex structure prediction using AlphaFold3^2^ and its clone model, Boltz-1^3^. Both prediction models utilize a “reference conformer” as one of their input features. When the chemical structure of the ligand is provided in SMILES notation or Chemical Component Dictionary (CCD) code, its three-dimensional structure (conformer) is generated using ETKDGv3 algorithm^7^ implemented in RDKit and then used as the reference conformer. This study systematically assessed how accurately the stereochemistry—specifically the chiralities, bond lengths, and bond angles—of the reference conformer is reproduced in the predicted structures.

To evaluate the stereochemistry of the ligands in the predicted structures, we utilized entries extracted from the PLINDER^8^ and PDBBind2020 dataset^9^. The criteria for selecting entries to construct the evaluation dataset were as follows:

1. Entries where proteins do not form multimers.
2. Entries containing one ligand that is not a polymer and thus can be represented by a single CCD code.
3. Entries with a ligand containing at least one chiral center at an sp3-hybridized carbon atom.
4. Entries where the non-hydrogen atoms of the ligand consist solely of the elements C, N, O, F, P, S, Cl, Br, and I.
5. Entries containing a ligand whose reference conformer can be generated by RDKit.

For the PLINDER dataset, we employed clustering based on a Tanimoto similarity threshold of 0.95 to exclude identical compounds. By applying these selection criteria, we extracted 6,600 and 4,500 entries from the PLINDER and PDBBind2020 datasets, respectively. The PLINDER dataset was further divided into “Before” and “After” based on whether the PDB entry’s release date is before or after the cut-off date of 2021-09-30. Thus, the “Before” split comprises structures included in the training data for AlphaFold3 and Boltz-1, while the “After” split serves as a test set comprising structures not used during model training. Although the PDBBind-based dataset shares many entries with PLINDER’s “Before” split, it also includes distinct compounds that are absent in the PLINDER dataset.

### Evaluation metrics

We created RDKit Mol objects from the SMILES notation found in the CCD and assigned conformers, including chirality, based on the three-dimensional structures of the compounds in the respective dataset. Using these RDKit Mol objects, we enumerated chiral centers and examined the R/S chirality of each atom according to the Cahn–Ingold– Prelog (CIP) priority rules implemented in RDKit^10^. We then assessed the chirality of the corresponding atoms in the predicted structures and evaluated their consistency. If even one atom exhibited a different chirality, we recorded it as inconsistent and evaluated the fraction of consistent molecules against the total entries. To simplify the problem, we focused solely on the chirality of *sp*^3^ carbon atoms, which are commonly found in biomolecules.

We further evaluated the deviation of the predicted structures from the reference conformer in terms of their bond lengths and angles. Specifically, we calculated the root mean square of the deviations (RMSDs) from the ideal bond lengths (𝑟̂*_i_*) and angles (𝜃*_i_*) of the reference conformers generated by ETKDGv3 algorithm of RDKit^7^.

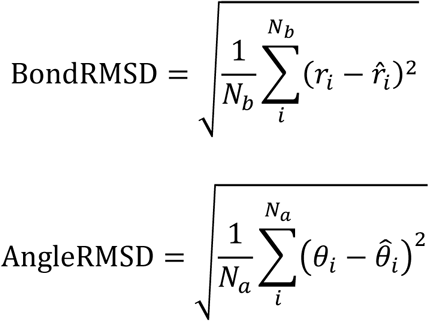

To assess the accuracy of ligand binding site predictions, we performed a least-squares superposition of protein residues within 10 Å of the ligand binding site and calculated the RMSD of the corresponding ligand atoms, as described in the AlphaFold3 paper^2^.

### Improvement of geometry through restraints

To overcome the stereochemistry violation problems described in the AlphaFold3 paper^2^, we developed a novel approach in this study. Instead of modifying or retraining the neural network model, we attempted to alter the diffusion model’s inference process to utilize chirality information, bond lengths, and bond angles as restraints. Our method was inspired by the symmetry-guided generation approach used in RFDiffusion^11^. In RFDiffusion, symmetric multimer protein structures are generated by applying hard constraints to protein Cα atoms at each step to enforce the desired symmetry. Similarly, we adopted an approach that applies restraints related to compound stereochemistry to the structures at each step of the reverse diffusion process. This method aims to accurately predict compound stereochemistry without requiring model retraining.

The diffusion model used in AlphaFold3 is the EDM^12^. In each inference step 𝑖 of EDM, the next step sample 𝒙_i+1_is calculated using the difference 𝒅_i_ computed from the output of the denoiser function 𝐷_θ_ and the current step sample 𝒙𝒙̂_i_:

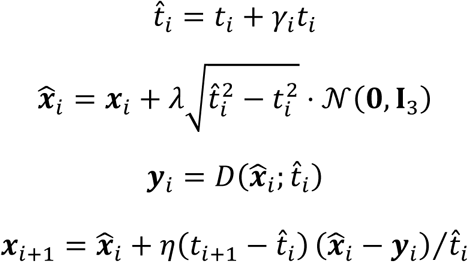

where, γ_i_, 𝜆, and 5 are hyperparameters to control the stochasticity levels.

To achieve denoising based on the training dataset, 𝐷 is implemented through a neural network 𝐷_θ_ with a parameter set 𝜃. This 𝒙_i+1_ is influenced by the noise scale and exhibits a distribution significantly different from the actual coordinate scale. This discrepancy makes applying restraints directly to 𝒙_i+1_impractical. In contrast, it was shown that the optimal denoising function 𝐷, which analytically minimizes the training objective, converges toward the mean of the dataset as the noise level increases. Conversely, as the noise level decreases, it approaches the ground truth of the dataset (see Equation B.3, eq. 57 in ref^12^). Consequently, the output of 𝐷_θ_is expected to remain consistent with the scale of the samples (i.e., experimental structures) present in the training dataset. Therefore, we implemented a method where stereochemical restraints based on the input reference conformer are applied to 𝒚_i_to obtain *ỹ_i_*, and then the difference between *x̂_i_* and 𝒚_i_ is used for calculating 𝒙_i+1_ for the subsequent step. These constrained reverse diffusion steps are applied when the condition 𝑡_i+1_ < 𝜎_start_ is satisfied, where 𝜎_start_ is an input parameter.

### Implementation of the restraints

In this study, we implemented stereochemical restraints targeting chirality of *sp*^3^ carbon atoms, bond lengths, and bond angles. These parameters typically do not deviate significantly from their ideal values calculated from the chemical structures when compounds are bound to the target protein molecules. In contrast, parameters such as dihedral angles can undergo substantial changes during protein-ligand binding processes and are therefore excluded from our restraint targets.

To maintain the correct chirality, we constrained the chiral volume to its ideal value. The chiral volume (𝑉_i_(𝑗, 𝑘, 𝑙)) for the chiral center atom (*i*) was calculated by the following formula:

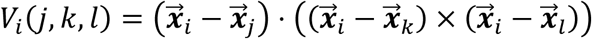

where the atoms *j*, *k*, *l* are adjucent atoms directly connected to atom *i* by covalent bonds. Then, the chirality restraints are achieved by minimizing the following term:

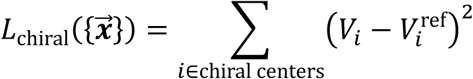

where 𝑉_i_represents the calculated chiral volume of the chiral center *i*, and 𝑉^ref^ is the reference value obtained from the input reference conformer. It is important to maintain the same order of *j*, *k*, and *l* in the above formula for both the current structure (𝑉_i_) and the reference conformer (𝑉^ref^) to preserve the same chirality.

For constraining bond lengths and bond angles within the compound, we minimized the following terms:

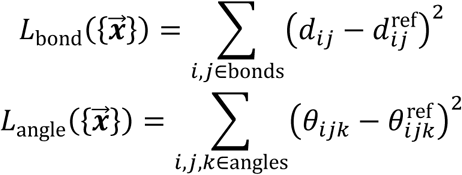

where 𝑑_ij_ denotes the bond length between atoms *i* and *j*, and 𝜃_ijk_ represents the bond angle formed by atoms *i*, *j*, and *k*. The reference values 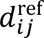 and 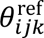 were derived from the input reference conformer.

The final restraint implementation was achieved by minimizing a weighted sum of the individual terms:

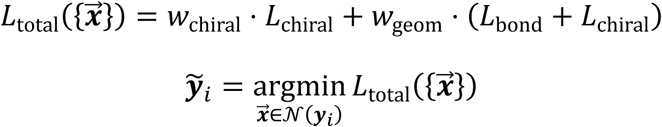

where 𝑤_chiral_and 𝑤_geom_ are weighting coefficients for each corresponding term. We implemented this method in Boltz-1^3^, a PyTorch-based clone of AlphaFold3. The minimization was performed using the conjugate gradient method implemented in the SciPy optimization library^13^.

### Multiple Sequence Alignment and Structure Prediction

Multiple sequence alignment (MSA) is a crucial component of the structure prediction pipeline of AlphaFold3 and Boltz-1, as evolutionary information significantly influences their structure prediction accuracy. In this study, we used the same pre-computed MSAs for all approaches, AlphaFold3, Boltz-1, and Boltz-1 with restraint-guided inference, to ensure fair comparison conditions. The MSAs were calculated using MMSeqs2 v15^14^ through colabfold_search implemented in LocalColabFold^15^. MMSeqs2 provides sufficient MSAs for most proteins while being significantly faster than the AlphaFold3’s pipeline that uses HMMER3^16,17^. For AlphaFold3 predictions, we used the default input parameters. For Boltz-1, to maintain consistency with AlphaFold3 conditions, we performed predictions with settings of recycle=10 and nsamples=5. All calculations were performed using NVIDIA GeForce RTX 4090 and H100 GPUs.

## Results

### Benchmark of protein-ligand complex structure prediction

In this study, we evaluated the protein-ligand complex prediction capabilities of AlphaFold3 (AF3)^2^ and Boltz-1^3^ using datasets derived from PLINDER^8^ and PDBbind2020^9^. Regarding the protein components of the predicted structures, both models exhibited excellent performance in structure prediction. For the PLINDER Before split, the Protein RMSD was ∼0.6 Å (Table 1), and similar values of ∼0.6 Å were obtained for the PDBbind dataset (Table S1). Even for the After split, which contains entries unseen by the models during training, both models achieved favorable performance with Protein RMSD values of ∼1.0 Å (Table 1, Figure 1A). Although performance declined in the After split compared to the Before split, both models consistently demonstrated excellent performance in protein structure prediction. In contrast, different trends were observed for the ligand structure prediction. For the PLINDER Before split, which consists of entries seen by the models, the Ligand RMSD was 3–4 Å (Figure 1B). Similarly, for the PDBbind dataset, also composed of entries seen by the models, the Ligand RMSD showed slightly lower RMSD values of ∼2.0 Å (Table S1). However, for the PLINDER After split, consisting of entries unseen by the models, Ligand RMSD deteriorated significantly, exceeding 7 Å (Figure 1B). These results are consistent with a previous report, which showed overfitting tendencies in AF3’s ligand binding pose predictions^18^.

**Figure 1.**
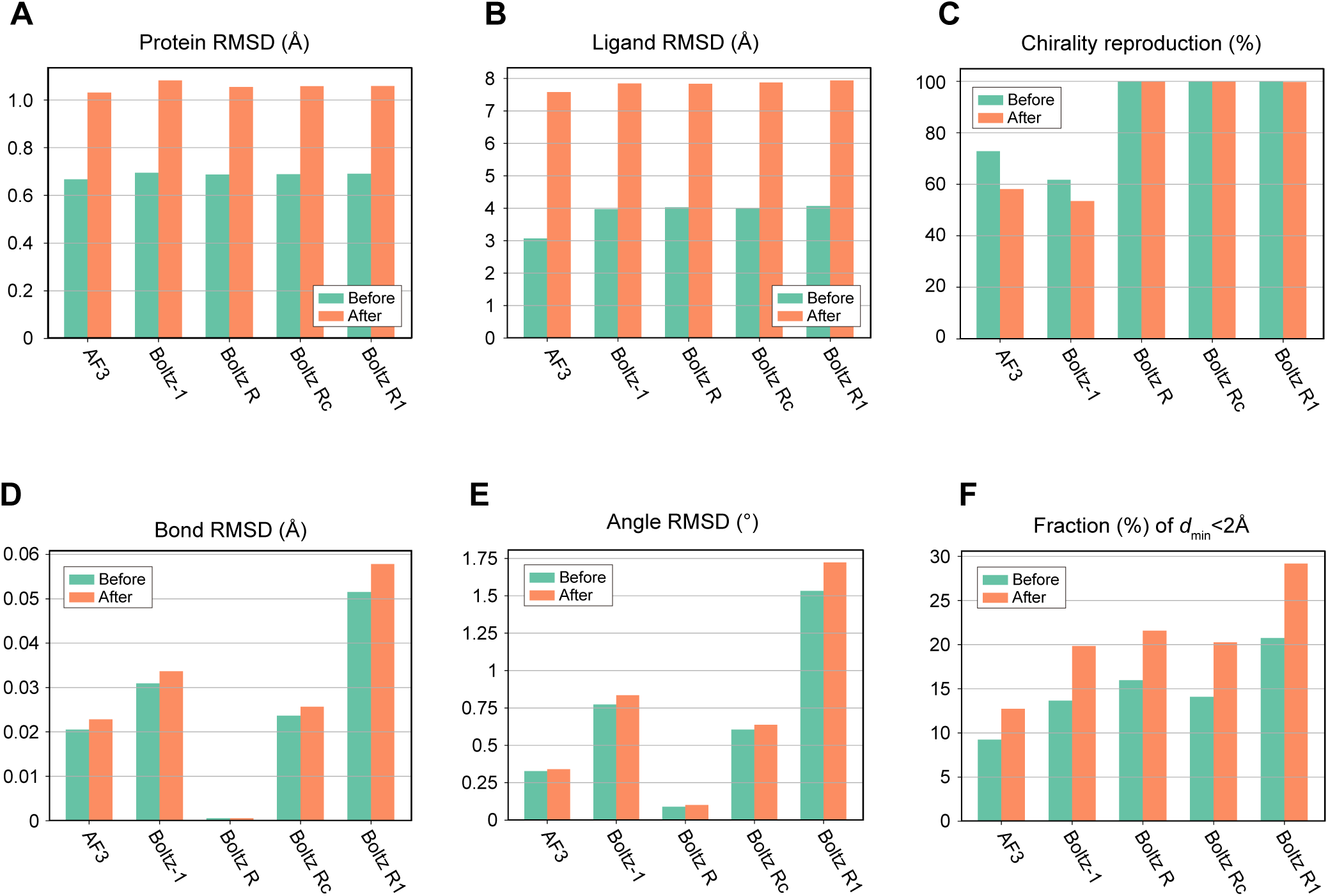
Performance evaluation of protein-ligand complex structure prediction methods on the PLINDER dataset. Bar charts comparing AlphaFold3 (AF3), Boltz-1, and Boltz-1 with restraint-guided inference variants (Boltz R, Boltz Rc, and Boltz R1). (A) Protein RMSD, (B) Ligand RMSD, (C) chirality reproduction rate, (D) Bond RMSD, (E) Angle RMSD, and (F) fraction of structures with steric clashes (*d*_min_ < 2 Å). Results are stratified by entry release dates: “Before” (green) represents entries released before the cut-off date (2021-09-30), while “After” (orange) represents subsequent entries.

**Table 1.**
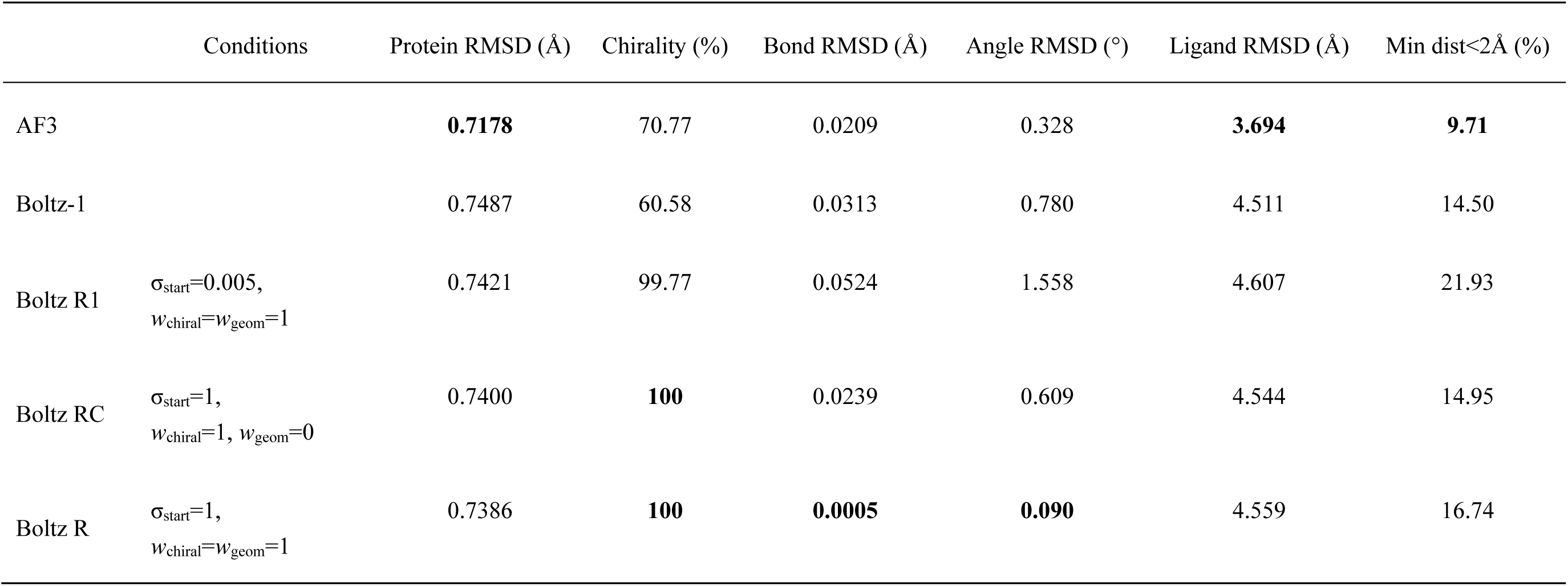
Evaluation metrics for proteins and ligands in the predicted structures from the PLINDER dataset used in this study.

Regarding the stereochemistry of the predicted ligand structures, we found an issue in reproducing the chirality of input molecules (Table 1, Figure 1C, Table S1). A substantial number of entries exhibited changes in chirality from the input structure. While the original AF3 paper reported error rates of only a few percent^2^, our dataset, which specifically focuses on chiral molecules, showed remarkably high error rates of 30–40%. One possible reason for this stereochemistry issue may stem from the ligand atom names used to create the input features. The atom names differ between the inputs generated using CCD code and those in SMILES notation. As a result, the input features can vary even for identical ligands, which might impact performance. To explore this hypothesis, we also compared the prediction results using CCD codes and found that chirality reproduction improved slightly (from 70.77% to 76.50% in PLINDER dataset). This aligns with the fact that the atom names from CCD are used during model training. However, a significant number of ligands still exhibited stereochemical errors.

A comparison of the results from AF3 and Boltz-1 revealed that AF3 showed superior performance in chirality reproduction, Bond, Angle, and Ligand RMSDs (Table 1, Figure 1). Overall, while AF3 generates higher-quality predicted structures than Boltz-1, both models still face challenges regarding ligand stereochemistry.

### Structure prediction using the restraint-guided inference

We also evaluated the prediction accuracy of Boltz-1 with the restraint-guided inference proposed in this study. This inference method involves multiple hyperparameters (𝜎_start_, 𝑤_chiral_, and 𝑤_geom_), so we assessed the model’s performance under several conditions (Table 1). Specifically, we conducted experiments where restraints were applied at the final step (𝜎_start_ = 0.005; Boltz R1) and during the last 55 steps (𝜎_start_ = 1; Boltz R) of the 200-step reverse diffusion process. Furthermore, we compared experiments that applied restraints only to the chiral volume (𝑤_chiral_ = 1 and 𝑤_geom_ = 0; Boltz Rc) against those that applied restraints to all parameters: chiral volume, bond length, and bond angle (𝑤_chiral_ = 𝑤_geom_ = 1; Boltz R).

The results showed that the chirality of the input molecule was reproduced 100% across all conditions with restraints except for Boltz R1, in which a small fraction of the chirality inversion was observed (Table 1, Figure 1C). These findings emphasize the importance of applying restraints iteratively within the reverse diffusion process. Additionally, results from Boltz Rc, which applied restraints solely on the chiral volumes, demonstrated that accurate chirality could be achieved by constraining only the chiral volume (Table 1, Figure 1C).

We then evaluated the Bond and Angle RMSD values for each prediction result. Under the Boltz R1 conditions, these values became worse compared to those from the model without restraints (Table 1, Figures 1D, 1E, 2A, 2B). This finding is consistent with the reproducibility of chirality, further showing the importance of applying multiple restraint steps. In contrast, under Boltz Rc conditions where restraints were applied only to the chiral volumes, the Bond and Angle RMSD values were comparable to those from the model without restraints (Table 1, Figures 1D, 1E, 2A, 2B). Thus, applying restraints on the chiral volumes does not adversely affect the bond length and angle geometries. One particularly notable finding was that under Boltz R conditions, where restraints were applied to all parameters (i.e., chiral volume, bond length, and angle), the Bond and Angle RMSD values showed significant improvement. They exceeded the values achieved by AF3 (Table 1, Figures 1D, 1E, 2A, 2B). This result again underscores that our method can accurately reproduce the stereochemical properties of the input reference conformers. Furthermore, we evaluated steric clashes between predicted protein and ligand structures by calculating the minimum distance (*d*_min_) between ligand atoms and protein atoms. We here defined a serious steric clash as the case where the *d*_min_ value fell below 2.0 Å, and evaluated the proportion of entries with such serious clashes over all entries.

In Boltz R1, where restraints were applied only to the final step of the reverse diffusion process, the highest percentage of serious steric clashes were observed (Table 1, Figure 1F). In contrast, in Boltz R and Boltz Rc, no significant increase in steric clashes was observed compared to Boltz-1 (Table 1, Figure 1F). These results show that multi-step restraints throughout the reverse diffusion process are necessary to prevent steric clashes.

### Case studies for chirality reproducibility of ligands

Next, we explored specific instances where the chirality of ligands differed from that in the input reference conformer. In the angiotensin-converting enzyme and inhibitor complex (Figure 3) (PDB ID: 4BXK)^19^, the chirality of one carbon atom was predicted to be different from that in the reference conformer (Figure 3C, D). We also analyzed the case where the ligand was specified by the CCD code. The resulting output structure also exhibited a different chirality from that in the input, exemplifying that the reproducibility of chirality was not improved using the atom name assignments derived from CCD (Figure 3E). In contrast, the Boltz R result with restraint-guided inference produced a structure with the correct chirality (Figure 3F). A comparison of the predicted structures showed that the Boltz-R result superposes better with the experimental structure than those by Boltz-1 or AF3 (Figure 3C, D, F). However, when measured by Ligand RMSD, AF3 and Boltz R exhibited similar values of 0.38 Å and 0.39 Å, respectively. The changes in the atomic positions due to the chirality change are not substantial, which are averaged out when evaluating RMSD. This result may explain why there were no significant changes in the distribution of Ligand RMSD despite improvements in chirality by the restraint-guided inference (Figure 2C).

**Figure 2.**
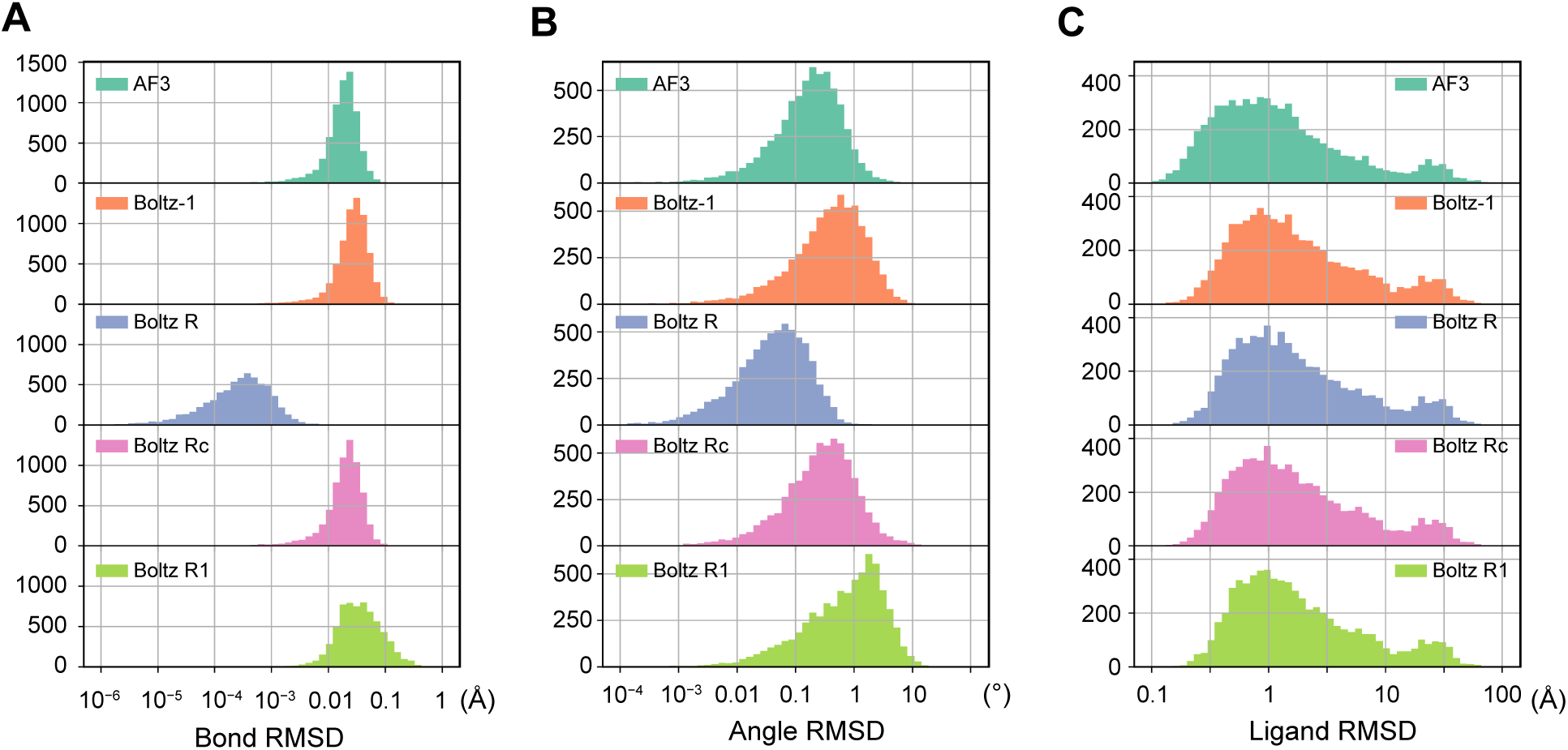
Distribution of ligand evaluation metrics on the PLINDER dataset. Histograms showing the distribution of prediction accuracy metrics for AlphaFold3 (AF3), Boltz-1, and Boltz-1 with restraint-guided inference variants (Boltz R, Boltz Rc, Boltz R1). (A) Bond RMSD, (B) Angle RMSD, and (C) Ligand RMSD distributions. All *x*-axes are displayed on logarithmic scales.

**Figure 3.**
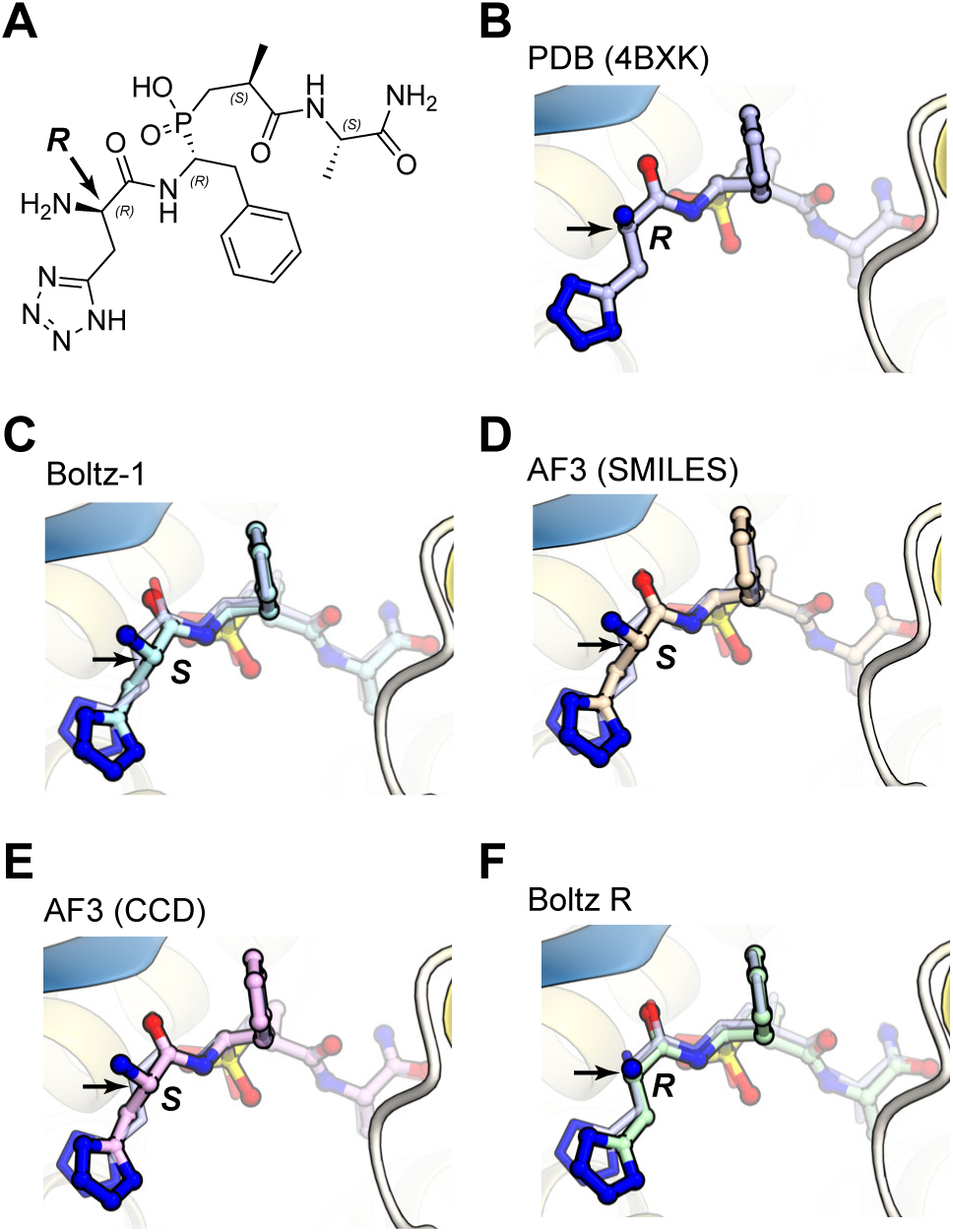
Chirality error in small molecule ligand prediction. Case study of angiotensin-converting enzyme and inhibitor complex (PDB ID: 4BXK). (A) Chemical structure of the ligand, (B) experimental structure, (C) Boltz-1 prediction, (D) AlphaFold3 prediction, (E) AlphaFold3 prediction using CCD code (1IU), and (F) Boltz-1 with restraint-guided inference (Boltz R). Proteins are depicted as ribbon models, and ligands are shown as ball-and-stick models. In panels (C)-(F), predicted ligand structures are overlaid with the semi-transparent experimental ligand structures for comparison. Arrows indicate chiral centers with stereochemical errors, and *R*/*S* designations denote their stereochemical configurations. Only the restraint-guided method (F) correctly reproduces the experimental chirality.

We also investigated the specific example of Cα carbon atom chirality in amino acid ligands. In the case of the glutamate transporter and D-aspartate complex (Figure 4) (PDB ID: 6R7R)^20^, both AF3 and Boltz-1 predicted conformations corresponding to the L-form amino acid (Figure 4C, D). Using CCD code to conform to the atom name nomenclature of amino acids (*i.e.*, CA, CB, etc.) did not improve the results (Figure 4E). In contrast, with the restraint-guided inference, Boltz-R predicted the structure with the correct chirality (Figure 4F). Similar results were also observed in other complex structures with D-form amino acids. These findings demonstrate that when the ligands resemble the monomers of biopolymers, such as L-amino acids in proteins or riboses in nucleic acids, AF3 and Boltz-1 tend to alter their chirality to match that of these biopolymers. While the input atom embeddings of AF3 include information on the atom names, elements, and formal charges, they lack specific information about the chiral centers. Consequently, the model cannot learn the relationship between the atom representations and the arrangement of substituents around the *sp*^3^-carbon atoms. Further examples demonstrating the chirality problem are shown in Figures S1–S4.

**Figure 4.**
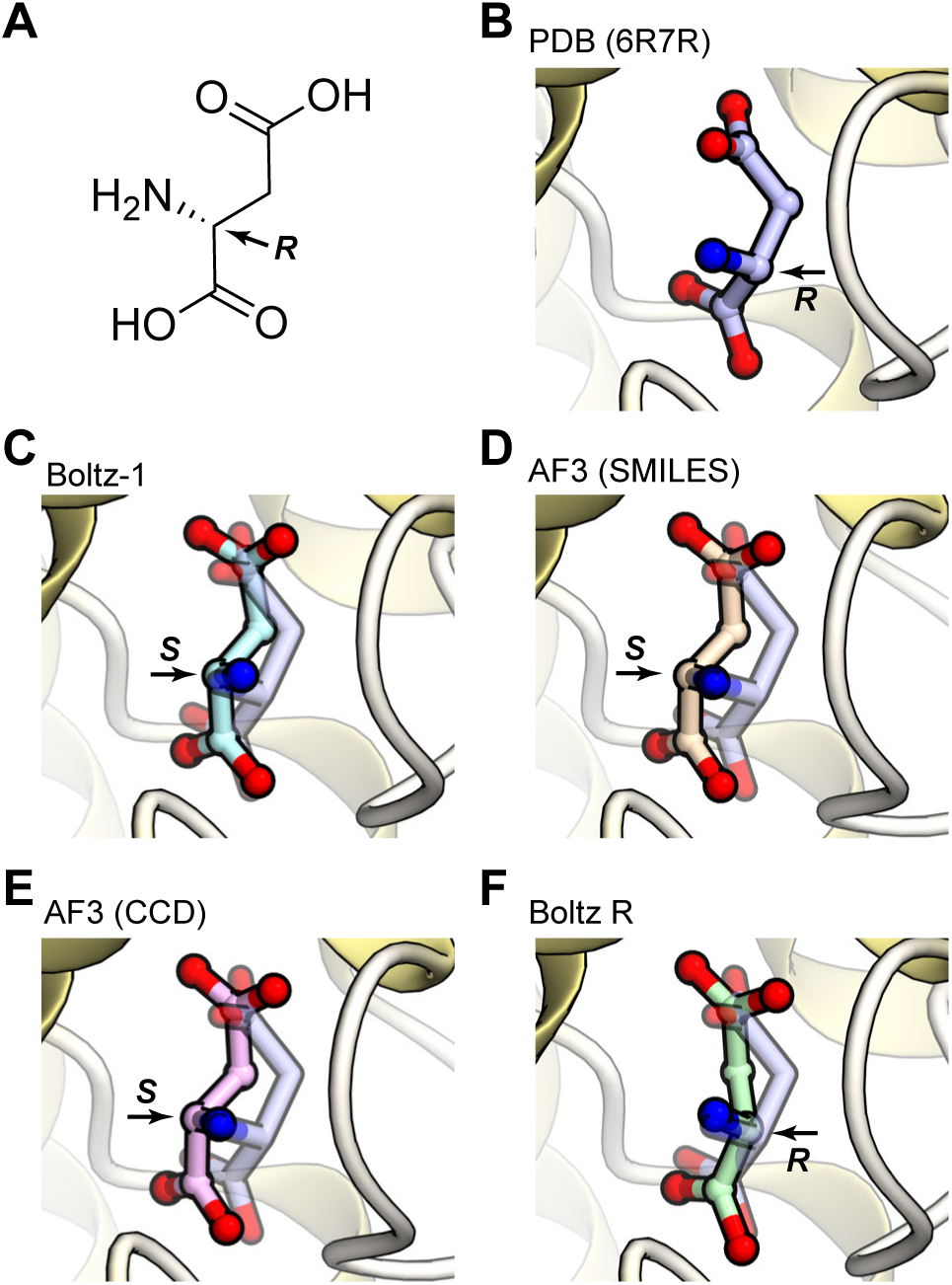
Chirality error in D-amino acid ligand prediction. Case study of glutamate transporter in complex with D-aspartate (PDB ID: 6R7R). (A) Chemical structure of the ligand, (B) experimental structure, (C) Boltz-1 prediction, (D) AlphaFold3 prediction, (E) AlphaFold3 prediction using CCD code (DAS), and (F) Boltz-1 with restraint-guided inference (Boltz R). Proteins are shown as ribbon models and ligands as ball-and-stick models. In panels (C)-(F), predicted ligand structures are overlaid with semi-transparent experimental ligand structure for comparison. Arrows indicate chiral centers with stereochemical errors, and *R*/*S* designations denote their stereochemical configurations. Only the restraint-guided method (F) correctly reproduces the chirality of D-aspartate.

### Case studies for bond and angle geometries of ligands

Next, we investigated specific cases where the bond and angle RMSDs of the ligand structures predicted by AF3 and Boltz-1 showed significant deterioration. In these cases, we observed several inconsistencies between the predicted ligand conformers and the reference conformers, including:

1. Structural inconsistencies in aromatic and non-aromatic rings
2. Hybridization state inconsistencies in non-cyclic structures

For the first case, we examined the ectoine-binding protein UehA complexed with ectoine (Figure 5) (PDB ID: 3FXB)^21^. In the structures predicted by AF3 and Boltz-1, the ligand’s ring structure was incorrectly modeled as planar, while the reference conformer contains a non-aromatic saturated ring (Figure 5). This error likely occurs because the chemical structure of the ring resembles that of pyrimidine nucleobase, which is commonly found in the training data. This similarity causes the model to predict it as an aromatic structure incorrectly (Figure 5D, E). For the second case, we examined the FimH lectin domain complexed with a small molecule (Figure 6) (PDB ID: 4AV4)^22^. In the prediction structure by AF3 and Boltz-1, the *sp*^1^ carbon atoms of the alkyne group in the ligand, which should form a linear structure, is instead predicted as an *sp*^3^ alkyl chain-like structure (Figure 6). This observation suggests that structures rarely found in natural biomolecules, such as *sp*^1^ carbons, are biased toward frequently occurring structures like *sp*³ carbons, resulting in predicted ligand structures with incorrect stereochemistry.

**Figure 5.**
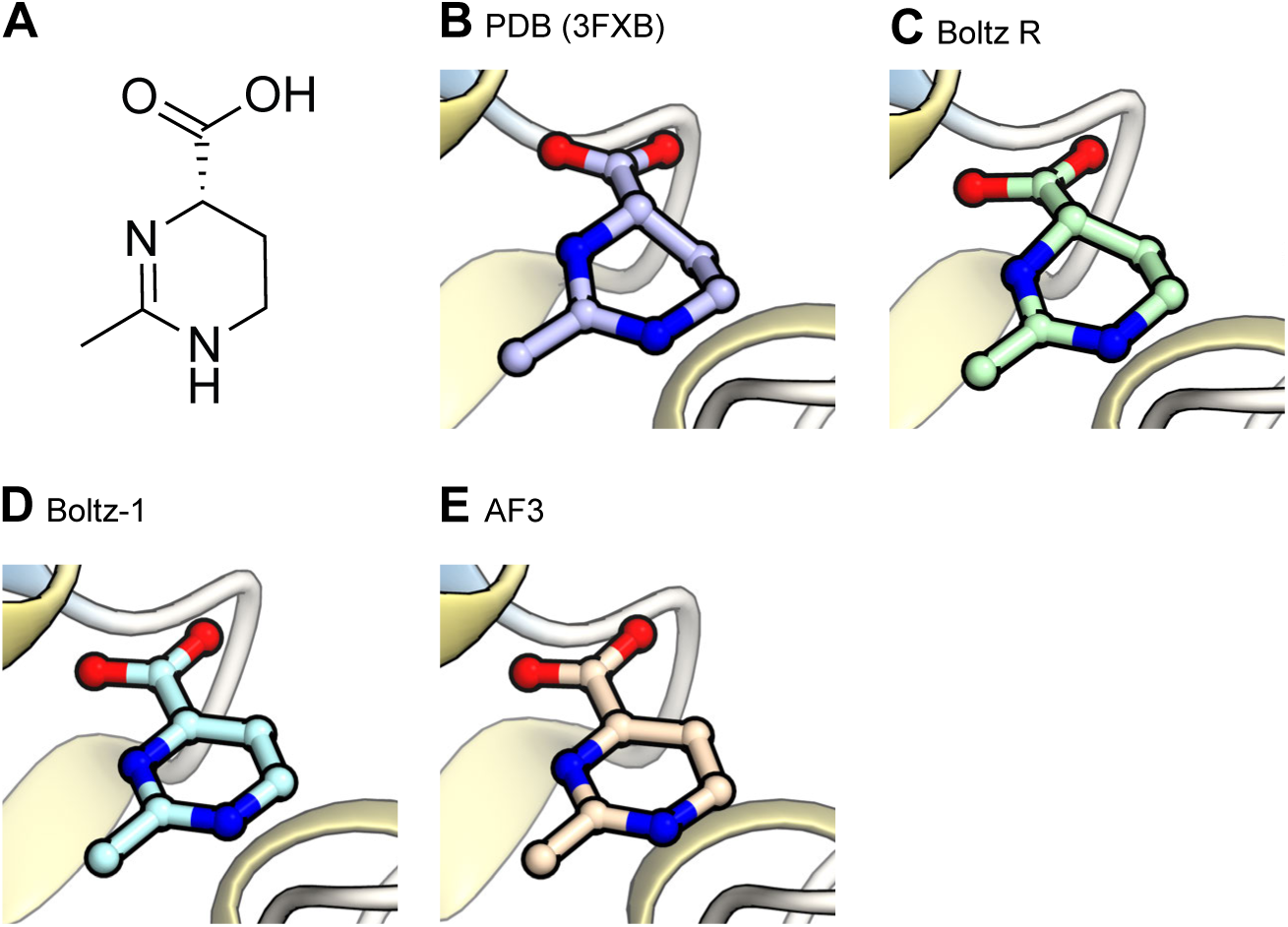
Ring conformation errors in ligand structure prediction. Case study of ectoine-binding protein UehA in complex with ectoine (PDB ID: 3FXB). (A) Chemical structure of the ligand, (B) experimental structure, (C) Boltz-1 with restraint-guided inference (Boltz R), (D) Boltz-1 prediction, and (E) AlphaFold3 prediction. Proteins are shown as ribbon models and ligands as ball-and-stick models. The ring structure is incorrectly predicted as planar in baseline methods (D, E), while the restraint-guided method (C) correctly reproduces the non-aromatic saturated ring conformation.

**Figure 6.**
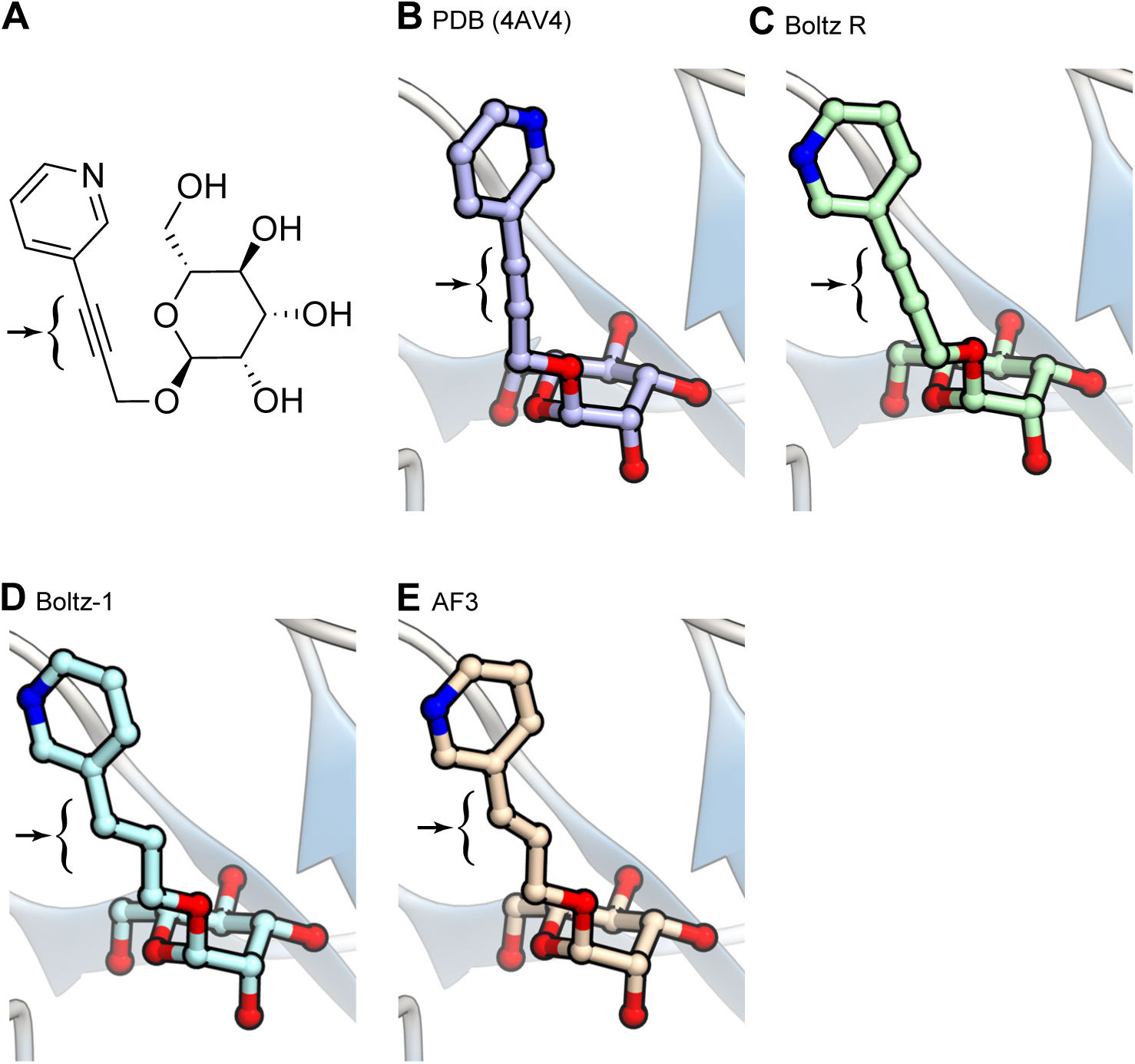
Bond and angle geometry errors in ligand structure prediction. Case study of FimH lectin domain in complex with a small molecule (PDB ID: 4AV4). (A) Chemical structure of the ligand, (B) experimental structure, (C) Boltz-1 with restraint-guided inference (Boltz R), (D) Boltz-1 prediction, and (E) AlphaFold3 prediction. Proteins are shown as ribbon models and ligands as ball-and-stick models. Arrows indicate the regions where the bond and angle geometry errors occur.

These problems may stem from fundamental limitations in the input feature representation of the AF3 model. Specifically, the atom and bond embeddings in AF3 do not include explicit information about hybridization states, as well as bond orders, such as single or double bonds. As a result, chemically distinct atoms and bonds are treated as similar input features. This leads the model to skew predictions toward structural patterns frequently occurring in the training dataset. The cases discussed here illustrate stereochemical errors likely caused by this input embedding issues. Even in such cases, our restraint-guided inference strategy successfully generated structures with correct stereochemical properties (Figures 5C, 6C). These results underscore that the proposed method effectively overcame the limitations of AF3 and Boltz-1 in ligand stereochemistry. Further examples demonstrating the bond angle geometry problem are shown in Figures S5 and S6.

## Discussion

In this study, we comprehensively investigated stereochemical accuracy in protein-ligand complex structure predictions by AlphaFold3 (AF3) and its clone model Boltz-1, using unified assessment criteria against the same benchmark dataset. Previous protein-ligand complex prediction methods, such as conventional molecular docking approaches^23^ and neural network-based methods like DiffDock^24^, typically treat dihedral angles as the target degrees of freedom for structure prediction. AlphaFold2 also uses dihedral angles as prediction targets for protein side-chain structures^1^. These methods can generate molecules with the desired stereochemical properties without problems. In contrast, AF3 employs a diffusion model approach and directly predicts the Cartesian coordinates of each atom in the target molecules. This methodology enabled the prediction of biomolecule and ligand structures within the unified and straightforward framework. However, the direct prediction of Cartesian coordinates creates a disadvantage in that it can generate molecules with incorrect stereochemistry. Our results confirmed that the ligands of the predicted structures contain significant levels of stereochemical errors in their chiralities, bond lengths, and angles. Then, we developed the restraint-guided inference method to address these stereochemical errors. Our evaluation confirmed that this method effectively improves the stereochemical validity of the ligand structures. Importantly, we demonstrated that the restraint-guided inference method does not adversely affect the performance of the binding pose prediction.

We focused on the method to address the stereochemical errors without retraining the model in this study. However, as discussed in the Results section, one potential cause of the ligand stereochemistry violations may arise from using feature representations, such as atom names, which can vary depending on the input notations. Therefore, a fundamental solution would require a model design that directly incorporates invariant features of compound structures, including hybridization states, chirality, and bond orders.

In the current models, natural biomolecules such as protein amino acids, nucleic acids, and glycans rarely experience those stereochemical issues. This is likely because these molecules are abundant in their training dataset. Therefore, a similar solution could theoretically be possible for small organic compounds by including sufficient numbers and diversity in the training dataset. However, the chemical space of small molecules is orders of magnitude larger than the limited set of biomolecular building blocks, making it impractical to train the neural network model to correctly preserve stereochemistry across arbitrary compounds. From this perspective, our approach provides a practical solution for the ligand stereochemistry problems.

## Author contributions

R.I. designed the study and analyzed the data. R.I., and Y.M. performed computational experiments and wrote the manuscript. R.I. directed and supervised this study.

## Acknowledgements

The authors thank Naruki Yoshikawa for constructive comments. This research was partially supported by the Research Support Project for Life Science and Drug Discovery (Basis for Supporting Innovative Drug Discovery and Life Science Research (BINDS)) from AMED under Grant Numbers JP25ama121027 for Y.M. and JP25ama121012 for R.I. This work was also supported by JSPS KAKENHI Grant-in-Aid for Transformative Research Areas (A) under Grant Numbers 25H01570 and 25H02250 for R.I. Additional support was provided by the Medical Research Center Initiative for High Depth Omics at the Institute of Science Tokyo, Nanken-Kyoten at the Institute of Science Tokyo 2025, and the Multilayered Stress Diseases project (JPMXP1323015483) at the Institute of Science Tokyo. This work utilized computational resources from the supercomputer “Flow” provided by the Information Technology Center at Nagoya University through the HPCI System Research Project (Project ID: hp240040) and the TSUBAME4.0 supercomputer at the Institute of Science Tokyo.

## Data and Software Availability

The materials supporting this article, including the source code and Colab notebook are available at github (https://github.com/cddlab/boltz_ext and https://github.com/cddlab/colabfold_boltz_restr/blob/main/Boltz1.ipynb).

