## Supplementary Information for "Improving Stereochemical Limitations in Protein-Ligand Complex Structure Prediction"

### Supplementary Figure Legends

#### Figure S1

Examples showing chirality errors in small molecule ligands.

Comparison of predicted ligand structures for (PDB ID: 8H59). (A) Chemical structure, (B) experimental structure, (C) Boltz-1 prediction with restraint-guided inference (Boltz R), (D) Boltz-1 prediction, and (E) AlphaFold3 prediction. Proteins are shown as ribbon models and ligands as ball-and-stick models. In panels (C)-(E), predicted ligand structures are overlaid with semi-transparent experimental ligand structure for comparison. Arrows indicate incorrectly predicted chiral centers with *R*/*S* designations indicating stereochemical configuration.

#### Figure S2

Examples showing chirality errors in small molecule ligands.

Comparison of predicted ligand structures for (PDB ID: 1ZRB). (A) Chemical structure, (B) experimental structure, (C) Boltz-1 prediction with restraint-guided inference (Boltz R), (D) Boltz-1 prediction, and (E) AlphaFold3 prediction. Proteins are shown as ribbon models and ligands as ball-and-stick models. In panels (C)-(E), predicted ligand structures are overlaid with semi-transparent experimental ligand structure for comparison. Arrows indicate incorrectly predicted chiral centers with *R*/*S* designations indicating stereochemical configuration.

#### Figure S3

Chirality errors in small molecule ligand prediction.

Comparison of predicted ligand structures for protein-ligand complex (PDB ID: 1OSV). (A) Experimental structure, (B) Boltz-1 prediction with restraint-guided inference (Boltz R), (C) Boltz-1 prediction, and (D) AlphaFold3 prediction. Proteins are shown as ribbon models and ligands as ball-and-stick models. In panels (B)-(D), predicted ligand structures are overlaid with semi-transparent experimental ligand structure for comparison. Each panel shows the chemical structure of the compound with *R*/*S* stereochemical notations at the chiral centers.

#### Figure S4

Chirality errors in small molecule ligand prediction.

Comparison of predicted ligand structures for protein-ligand complex (PDB ID: 5Z2C). (A) Experimental structure, (B) Boltz-1 prediction with restraint-guided inference (Boltz R), (C) Boltz-1 prediction, and (D) AlphaFold3 prediction. Proteins are shown as ribbon models and ligands as ball-and-stick models. In panels (B)-(D), predicted ligand structures are overlaid with semi-transparent experimental ligand structure for comparison. Each panel shows the chemical structure of the compound with *R*/*S* stereochemical notations at the chiral centers.

#### Figure S5

An Example showing errors in the ligand bond and angle geometries.

Comparison of predicted ligand structures for (PDB ID: 5IWE). (A) Chemical structure, (B) experimental structure, (C) Boltz-1 prediction with restraint-guided inference (Boltz R), (D) Boltz-1 prediction, and (E) AlphaFold3 prediction. Proteins are shown as ribbon models and ligands as ball-and-stick models.

#### Figure S6

An Example showing errors in the ligand bond and angle geometries.

Comparison of predicted ligand structures for (PDB ID: 7XK9). (A) Chemical structure, (B) experimental structure, (C) Boltz-1 prediction with restraint-guided inference (Boltz R), (D) Boltz-1 prediction, and (E) AlphaFold3 prediction. Proteins are shown as ribbon models and ligands as ball-and-stick models.

#### Table S1

Evaluation metrics for proteins and ligands in the predicted structures from the PDBBind dataset used in this study.

| | Conditions | Protein RMSD (Å) | Chirality (%) | Bond RMSD (Å) | Angle RMSD (°) | Ligand RMSD (Å) |
|---|---|---|---|---|---|---|
| AF3 | | 0.660 | 78.6 | 0.0196 | 0.244 | 1.871 |
| Boltz-1 | | 0.629 | 67.6 | 0.0303 | 0.650 | 2.534 |
| Boltz R1 | $\sigma_{\text{start}} = 0.005, w_{\text{chiral}} = w_{\text{geom}} = 1$ | 0.624 | 99.9 | 0.0494 | 1.306 | 2.550 |
| Boltz Rc | $\sigma_{\text{start}} = 1, w_{\text{chiral}} = 1, w_{\text{geom}} = 0$ | 0.628 | 100 | 0.0234 | 0.509 | 2.537 |
| Boltz R | $\sigma_{\text{start}} = 1, w_{\text{chiral}} = w_{\text{geom}} = 1$ | 0.628 | 100 | 0.0005 | 0.044 | 2.516 |

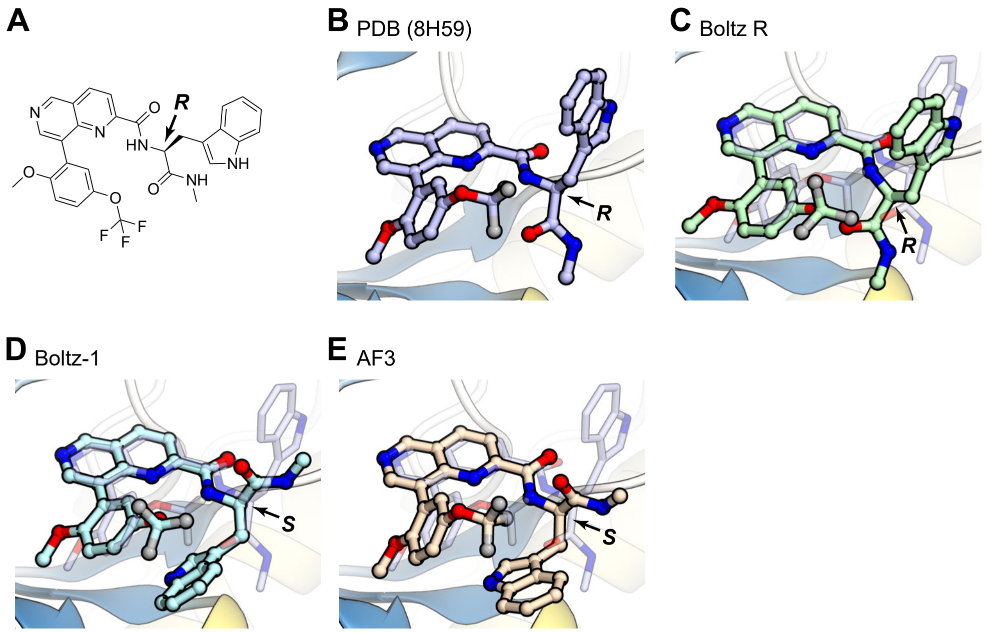


### Supplementary Figure S1

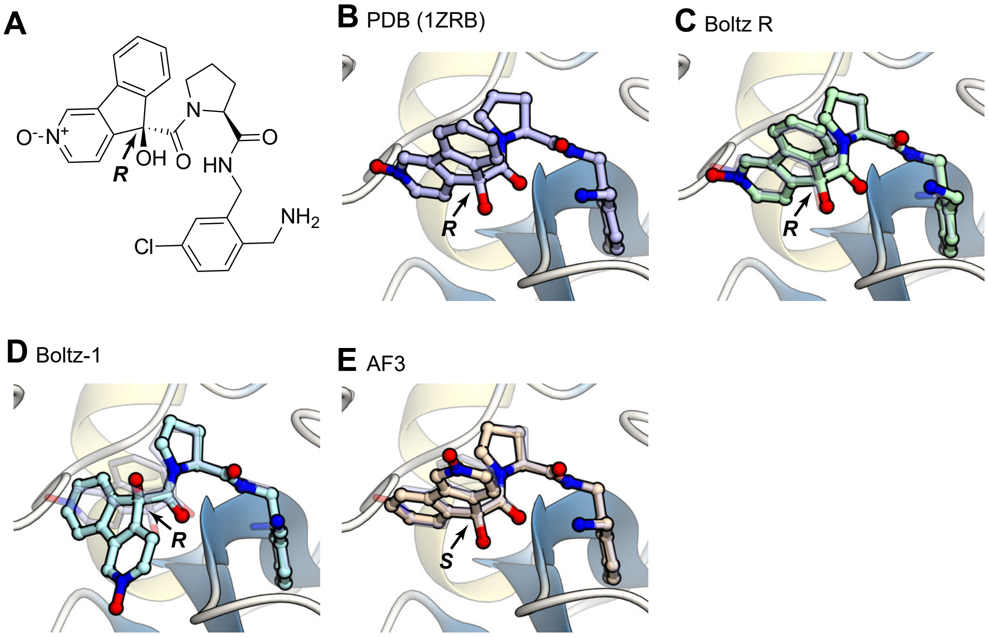


**Supplementary Figure S2**

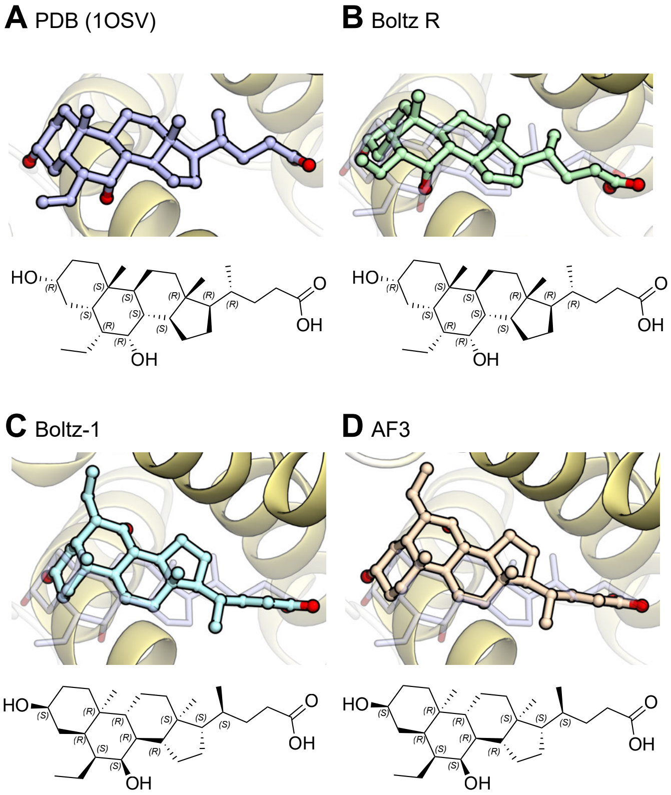


**Supplementary Figure S3**

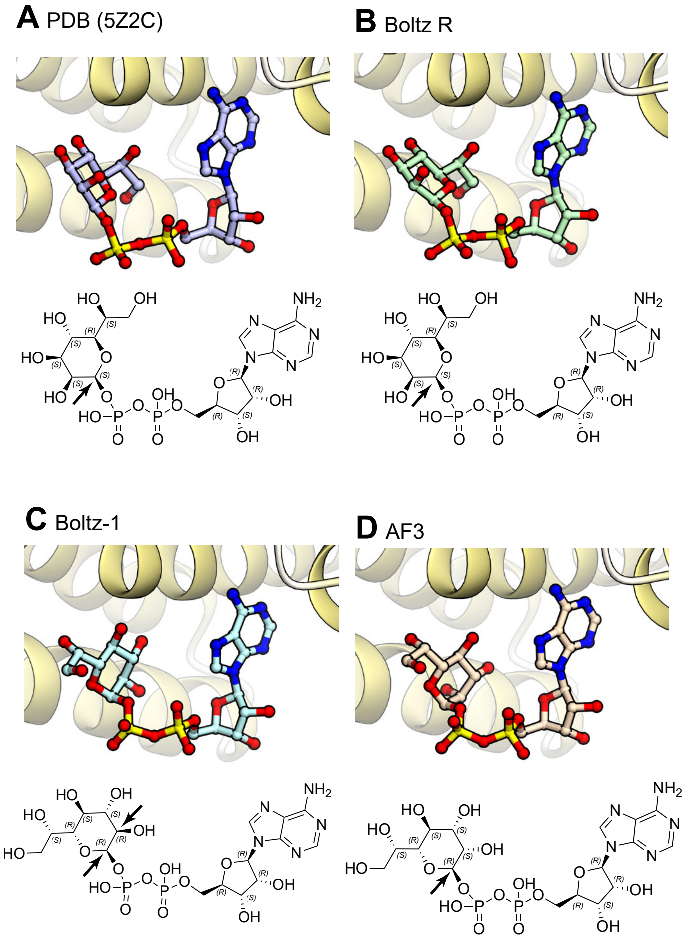


**Supplementary Figure S4**

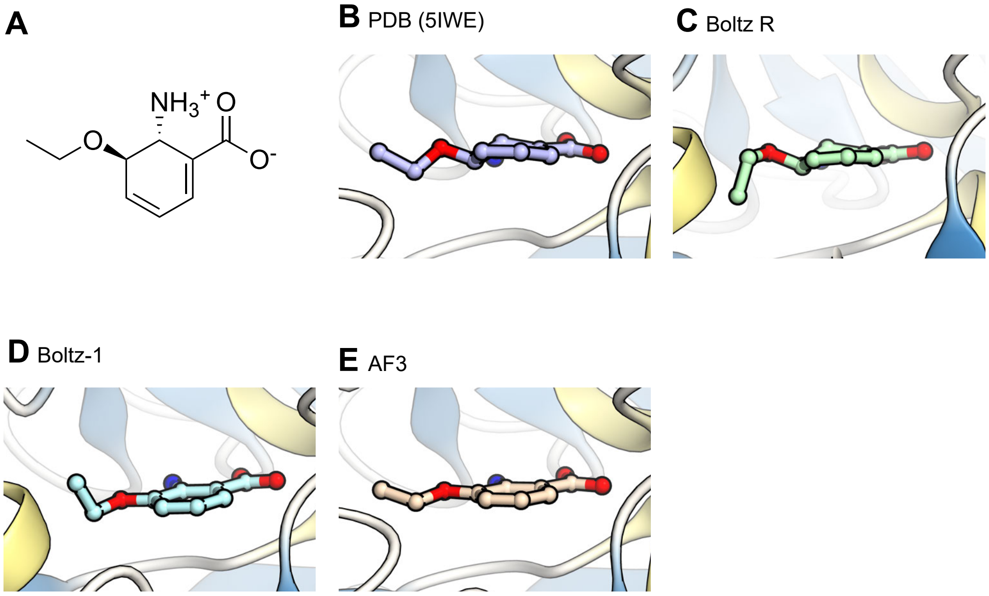


**Supplementary Figure S5**

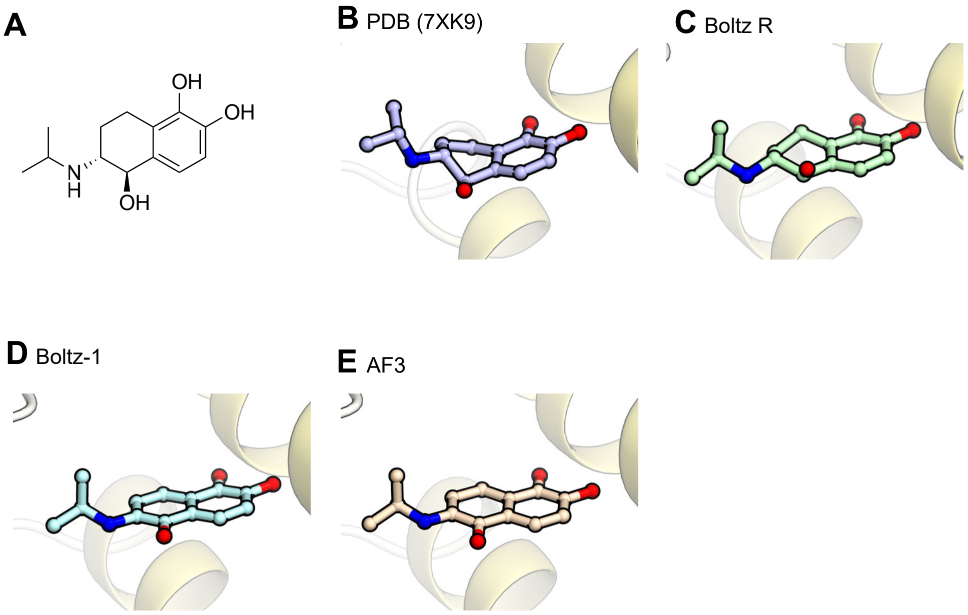


**Supplementary Figure S6**
